# Scaling Precision Medicine: Singapore’s PRECISE-SG100K Genomic Engine

**DOI:** 10.1101/2025.03.13.642552

**Authors:** Claire Bellis, Gabriel Kolle, Jacklyn Yong, Maxime Hebrard, Bitong Clarabelle Alexandrine Lin, Tat Hung Koh, Paul CP Cheng, Shakita D/O Sunmugam, Khai Koon Heng, Zhicheng Xie, Wee Yang Meah, Xiao Yin Chen, Erin Yi Ting Lim, Jyn Ling Kuan, Rudi Alberts, Helen Speirs, Sherilyn Lim, Yen Lynn Wong, Siew Hong Leong, Jack Ling Ow, Rodrigo Toro Jimenez, Eleanor Wong, Terry Yoke Yin Tong, Swat Kim Kerk, Jiali Yao, Miao Ling Chee, PRECISE-SG100K Genomic Engine Consortium, Khung Keong Yeo, Ching-Yu Cheng, Xueling Sim, Weiling Zheng, Chiea Chuen Khor, Shih Wee Seow, E Shyong Tai, John C Chambers, Nicolas Bertin, Patrick Tan

## Abstract

The whole-of-nation National Precision Medicine (NPM) programme in Singapore aims to integrate a genomics backbone into standard healthcare. Phase II recently completed the sequencing of 102,202 genomes from the diverse Asian population of Singapore, generating the PRECISE-SG100K dataset. The scaling from the pilot SG10K_Health phase to national production scale demanded a central shift: from earlier generation workflows to an integrated, industrial grade “***Genomic Engine***”. The initiative was co-developed under a public-private partnership model. Our Genomic Engine was designed with an automation-first approach, as a modular platform linking a purpose-built, centralized biospecimen processing and DNA extraction facility (EXTRACT Lab), a high-throughput sequencing operation (Illumina Data Generation Infrastructure, IDGI), with the cloud-native analytics (GRIDS) running on a fully integrated PRECISE Sample Tracking System (STS).

Here we describe the key design features of our Genomic Engine. Highlighting the Stress Test phase where the automated DNA extraction yields failed to meet expectations. We share the operational adjustments which were implemented to stabilize production and resulted in a three-fold reduction in sample triage due to improved DNA recovery. Without establishing the operational guardrails during the Stress Test, the Engine would not have reached a steady-state production evidenced by >3,000 genomes per month, median turnaround time 11 days, 99.6% operational success, within in six months. This framework offers a practical blueprint for other population-scale initiatives seeking to bridge the gap between scientific piloting and industrial delivery.

## Main text

Precision medicine holds immense promise in transforming healthcare systems by leveraging genomic insights to inform individualized care. Singapore has taken a national approach to this transformation through its National Precision Medicine (NPM) program, initiated in 2017. As a starting proof of concept^1, 2^, NPM successfully sequenced 10,000 participants from Singapore’s multi-ethnic population (Supplementary Note 2: Singapore National Precision Medicine Program – Background). The second phase, PRECISE-SG100K, represents the program’s proof of value and has recently completed sequencing 100,000 whole genomes from Singapore’s major longitudinal and population cohort studies (Figure 1 Box A; Supplementary Note 1: Singapore National Precision Medicine Program Phase I – Lessons Learned), with the ultimate goal of integrating genomic data into clinical workflows.

**Figure 1.**
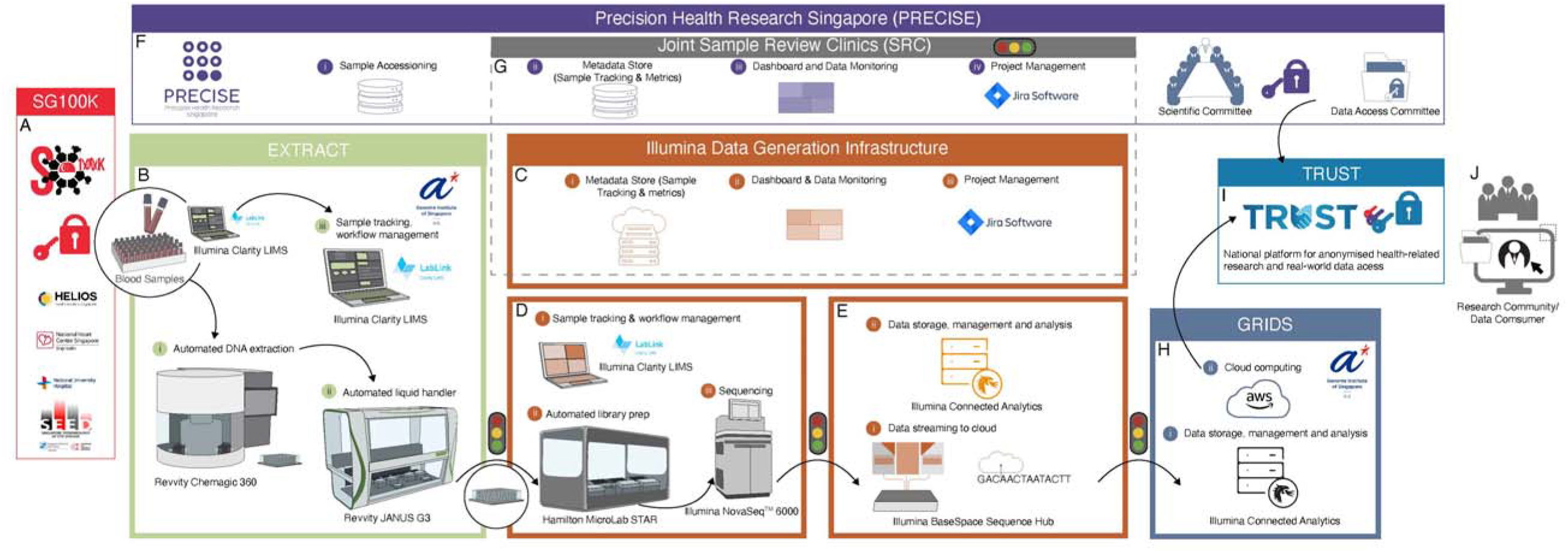
PRECISE-SG100K Genomic Engine Architecture consisted of automated workflows with minimized user interventions. **<u>Box A:</u>** The SG100K population and longitudinal studies contributing whole-blood biospecimens for NPM Phase II. Cohort sites prepare pre-barcoded 96-well plates (500µl each) with digital manifests submitted via Illumina Clarity LIMS LabLink. Plates are inspected, sealed, and transferred under chain-of-custody to the centralized EXTRACT facility**; <u>Box B:</u>** Centralized high-throughput facility for automated DNA extraction, quantification, and normalization from SG100K biospecimens. Operates under harmonized SOPs to attain a production-scale capacity of >1,500 sequencing-ready samples per week with consistent, high-quality yield and purity; **<u>Box C-E:</u>** Illumina Data Generation Infrastructure (IDGI) is a scalable Whole Genome Sequencing (WGS) data generation module that enables > 800 sample throughput per week starting from DNA plates and completing with delivery of high-quality data. IDGI consists of automated Library Preparation and Sequencing (Box D), a cloud-based DNA analysis module leveraging BaseSpace™ Sequence Hub, Illumina Connected Analytics (ICA) and DRAGEN™ that manages demultiplexing, secondary analysis and data delivery (Box E) which is overlayed by a sample management system which enables process tracking, monitoring and visibility across the entire workflow (Box C). Submission of sample data from Extract (Box B) and delivery of data to GRIDS (Box F) are fully automated processes; **<u>Box F:</u>** GRIDS built the PRECISE Sample Tracking System (STS) with the goal of recording and monitoring metadata, status, and quality metrics associated with each sample flowing between the different modules of the PRECISE-SG100K Genomic Engine. The STS communicated with each of the parties and centralized relevant information from each process or “handshake”. End-to-end digital infrastructure linking all modules across the Genomic Engine. Integrates Clarity LIMS, AWS DynamoDB, and SNS-Lambda notifications to provide real-time visibility of sample status and metadata. Enables traceability, automated alerts, and centralized dashboards for operational monitoring and auditability; **<u>Box G:</u>** Building upon the data flow established in Box C, biweekly sample data was aggregated and presented to key stakeholders during Joint Sample Review Clinics. This collaborative process was supported by the streamlined transfer of data from ICA to Jira, enabling efficient tracking and resolution of samples requiring further assessment or failing to meet contractual standards; **<u>Box H:</u>** Once high-quality genomic data was deemed ready to deliver to PRECISE, IDGI sent a notification GRIDS Module. A workflow, running within ICA, copied the data from ILMN managed storage to PRECISE managed storage ensuring data handover and ownership. Once the transfer was completed, a notification was sent to Illumina. In case of transfer failure, GRIDS and ILMN teams investigated the root cause, then transfer was triggered manually. Throughout this process, the genomic data is encrypted in transit and at rest. Once in PRECISE managed storage, lifecycle rules transitioned the data into long term archive. Processes related to DNA sequencing, mapping and variant calling were initially recorded by ILMN in ICA. The relevant information was then shared with PRECISE. Notifications related to data delivery were initially recorded by the Data Management and Repository Module. The underlying database was shared with PRECISE STS system.Leveraging highly distributed compute, GRIDS process single sample WGS data through standardized pipelines for joint variant calling, quality control, data releases and variant annotation **<u>Box I:</u>** Data releases are transferred to TRUST, a secure platform manage by the Ministry of Health. Genomic data is de-identified and linked to clinical data and electronic medical records into the highly controlled TRUST environment**; <u>Box J:</u>** Access to the PRECISE-SG100K data release by the researcher community is managed by a data access committee that review applicant identity and project goals prior to enable secured access to the data through the TRUST platform.

The first phase, SG10K_Health was scientifically productive, generating a multiethnic reference panel^2^ and invaluable insight into the prevalence on clinically actionable genetic variation^3^ but faced scaling issues due to distributed workflows and manual processes (Supplementary Note 1: Singapore National Precision Medicine Program Phase I – Lessons Learned). Excessive variability in DNA quality, fragmented sequencing operations, and a lack of standardized metadata protocols led to inefficiencies and constrained reproducibility. These challenges drove the conceptualization and implementation of an integrated Genomic Engine for the second phase, designed to support scale, traceability, and throughput by emphasising an automation-first principle.

Public-private collaboration was key in the second phase, with partnerships between PRECISE, the Genome Institute of Singapore, Illumina, and a local sequencing service provider co-developing and operating the Genomics Engine. This collaboration facilitated access to commercial-grade technologies, minimized infrastructure duplication, and promoted knowledge transfer. The co-location of industry and research teams allowed for real-time troubleshooting and continuous process improvement, essential for creating a system capable of national-scale delivery.

Industry partners played critical roles in capacity planning, logistics optimization, and technology training, allowing Singapore to achieve operational scale quickly. Formalized frameworks for cost-sharing and IP management provided alignment across public health goals and commercial interests with stakeholders engaged in joint execution and shared responsibility for delivery of data.

The resulting PRECISE-SG100K Genomic Engine, developed through this public-private partnership combined academic insight with commercial expertise, offering a fully integrated architecture for comprehensive genome processing. This system includes modular components for centralized sample tracking, DNA extraction, automated sequencing, and cloud-based informatics (Figure 1).

The EXTRACT DNA laboratory forms the intake point of the Genomic Engine (Figure 1 Box B; Supplementary Note 3: EXTRACT Lab Module – Workflow, Biospecimens, and Quality Control). As a high-throughput, centralized facility, it employs a high level of automation to optimise and standardize extraction from blood-based biospecimens, ensuring consistent DNA quality and concentration. Operating under harmonized, standard operating procedures, the lab eliminated much of the sample-to-sample variability observed in the pilot phase and has ensured operational consistency in Phase II, with the capacity of generating >1,500 sequencing ready DNA samples on a weekly basis.

Library preparation, sequencing and secondary analysis were driven by the Illumina Data Generation Infrastructure (Figure 1 Boxes C-E; Supplementary Note 4. Illumina Data Generation Infrastructure). This modular architecture comprises an automated, laboratory setup with production scale sequencing instruments that feed into a high throughput data production environment for primary and secondary data analysis. Specifically, DRAGEN™ germline pipeline version 3.7 was adopted by the program in alignment with other large-scale population genome sequencing initiatives^4, 5^. The infrastructure enabled steady-state data generation and delivery of >800 genomes per week, with consistent depth and quality metrics.

Informatics and data stewardship are managed by the Genome Research Informatics & Data Science (GRIDS) platform which deployed a secure environment for the processing, storage, and analysis of genomic data. GRIDS bridges the data production and downstream analysis by harmonizing outputs and providing standardized pipelines for joint variant calling, quality control, data releases and interpretation-ready variant annotation (Figure 1 Box H; Supplementary Note 5. Genome Research Informatics & Data Science - Informatics and data stewardship). The platform enforces data governance and access frameworks aligned with both national and international standards^6^, and enables the transfer of high-quality, functionally annotated variant catalogues at key release stages (10K, 50K, 100K) to TRUST (Figure 1 Box I; Supplementary Note 5. Genome Research Informatics & Data Science - Informatics and data stewardship), a secure national platform that integrates genomic data with participants’ Electronic Medical Records as well as health and wellness information collected by PRECISE-SG100K longitudinal and population cohorts.

Sample-level provenance, and data auditability were maintained through a Laboratory Information Management System, tailored to project-specific workflows (Figure 1 Box F; Supplementary Note 4. Illumina Data Generation Infrastructure) and complemented by purpose-built web-services for tracking of sample status from collection to analysis, linking to associated metadata, run performance metrics, and status logs. Operational transparency was maintained by establishing bi-weekly Sample Review Clinics, in which all key stakeholders convened to monitor progress, identify early deviation in data quality and arbiter decisions on exception handling including re-extraction, sequencing top-up, potential sample swaps, or contamination (Figure 1 Box F; Supplementary Note 6: Operational Considerations in the PRECISE-SG100K Genomic Engine), ensuring data integrity for use for research and future downstream clinical applications.

The PRECISE-SG100K Genomic Engine delivered 102,202 genomes in just over 3 years. 99.6% of samples met or exceeded all quality metrics set by the program including coverage, yield, variant callability, cross-contamination. The median turnround time from sample receipt to delivery of a high-quality genomic data bundle was 11 days (Supplementary Note 4. Illumina Data Generation Infrastructure, Table 6).

Designed for extensibility, the engine can handle various sample types, including archived DNA and clotted blood and supports flexible pipeline adaptation for transcriptomic, and epigenomic analyses. These capabilities provide a future-facing framework for evolving clinical and research needs, including the integration of diverse sequencing platforms, multi-omics data modalities and emerging data analytics technologies^7^.

Singapore’s PRECISE-SG100K Genomic Engine offers a model for delivering precision medicine infrastructure at national scale. Its automation-forward architecture and deeply integrated public-private partnership provide a blueprint for institutions and countries seeking to develop population-wide genomics initiatives. As healthcare shifts toward individualized care, such scalable Genomic Engines will be central to realising the promise of genomic medicine. By detailing the “how” as much as the “what”, we hope this correspondence will provide the granular operational insights and practical guidance for policymakers, health systems, and large population scale precision medicine programmes seeking to integrate genomic insights to transform healthcare systems.

## Supporting information

Standalone Supplemental Notes referenced in Summary Manuscript

## Data availability

The PRECISE-SG100K whole genome sequence data described in this manuscript is under controlled access due to local regulations.

Data access request can be submitted to the NPM Phase II / PRECISE-SG100K Data Access Committee by emailing PRECISE for details.

## Code availability

Code used during the current study are available from the corresponding author on reasonable request.

## Acknowledgements

We thank all participants, the study team, and the investigators for their research contributions. The authors acknowledge the funding support from the Genome Institute of Singapore, Agency for Science, Technology and Research (A*STAR). The Saw Swee Hock School of Public Health Tissue Repository and National University Hospital Tissue Repository provided long-term storage, retrieval, and preparation of SPHS and SERI biospecimens. We would like to acknowledge the support of and thank the Genomic Engine Steering Committee, staff of PRECISE, and GIS sequencing platform and Office of Procurement. NHCS Biobank provided long-term storage, retrieval, and preparation of SingHEART cohort biospecimens. We also thank Professor James Thompson and Mr William Day, Karolinska Institute Biobank, Jeffrey G Meyer, Mayo Clinic, All of Us, and Drs Jia Kia Lim and Uwe Jaentges from Revvity for DNA extraction optimization guidance and advice.

## Author contributions

The manuscript was drafted by C.B., G.K., J.Y., and M.H., with support from R.A., N.B., S.L., L.S.H., E.W., W.Z., and P.T. edited by P.T., and jointly revised and approved by all co-authors. C.B. coordinated manuscript submission-related administrative tasks. J.Y. conceived the concept, designed, and prepared the figure and table. B.C.A.L., Z.X., T.H.K., P.C.P.C., S.S., K.K.H., Z.X., W.Y.M., X.Y.C., E.Y.T.L. optimized and operationalized EXTRACT DNA processing workflow. C.B., B.C.A.L., Z.X., X.Y.C., E.Y.T.L., RA., N.B., M.H., J.O., R.T.J. curated data. C.B., B.C.A.L., Z.X., X.Y.C., E.Y.T.L., M.H., J.L.O., N.B., R.T.J., R.A. performed the data analyses. M.H., J.O., N.B., R.T.J. built the PRECISE sample tracking system. T.Y.Y.T., S.K.K., J.Y., and M.L.C. coordinated SG100K population study cohorts. C.C.K. is principal investigator of the EXTRACT laboratory. K.K.Y., C.Y.C., X.S., J.C.C. are principal investigators for the SG100K cohorts. Members of the *PRECISE-SG100K Genomic Engine Consortium* community are listed in alphabetical order. P.T., J.C.C., and E.S.T. conceived and designed Singapore’s National Precision Medicine programme.

## Funding and IRB Statements

This research is supported by the National Research Foundation Singapore under NPM program Phase II funding (MOH-000588) administered by the Singapore MOH’s National Medical Research Council. The NPM Phase I (SG10K_Health) project was funded by the Industry Alignment Fund (Pre-Positioning) (IAF-PP: H17/01/a0/007). We acknowledge the support of GIS/A*STAR core funding. We express our thanks to participants of the HELIOS study and the HELIOS operation team for recruitment, organisation, and data/sample collection. This study (NTU IRB: 2016-11-030) is supported by is supported by Singapore Ministry of Health’s (MOH) National Medical Research Council (NMRC) under its OF-LCG funding scheme (MOH-000271-00), the National Research Foundation, Singapore, through the Singapore Ministry of Health’s National Medical Research Council and the Precision Health Research, Singapore (PRECISE) under the National Precision Medicine programme and intramural funding from Nanyang Technological University, Lee Kong Chian School of Medicine and the National Healthcare Group. The SPHS studies are supported by award schemes from the Biomedical Research Council (BMRC) of Singapore, National Medical Research Council (NMRC) of Singapore [including Large Collaborative Grant MOH-000271-00], and infrastructure funding from the Singapore Ministry of Health (Population Health Metrics and Analytics PHMA), National University of Singapore and National University Health System, Singapore. SingHeart study received grant support in memory of Mr Henry H L Kwee as well as from Lee foundation. This work was also supported by core funding from SingHealth and Duke-NUS Institute of Precision Medicine (PRISM) and centre grant awarded to the National Heart Centre Singapore from the National Medical Research Council, Ministry of Health, Singapore (NMRC/CG/M006/2017_NHCS and MOH-000985). The SERI study is supported by Singapore’s MOH NMRC under its CS-IRG (NMRC/CIRG/1488/2018) and CSA (MOH-CSASI22jul-0001), and OF-IRG (NMRC/OFLCG/004/2018) funding scheme. We express our thanks to participants of the Singapore Epidemiology of Eye Diseases (SEED) study. This study was approved by individual institutional review board (IRB) study protocol approvals, and EXTRACT obtained requisite clearance from full application by the Human Biomedical Research Office, A*STAR, IRB (IRB Reference: 2021-191).

## PRECISE-SG100K Genomic Engine Consortium

Shimin Ang^6^, Graeme Bethel^4^, Daisy Chae^5^, Miao Li Chee^14^, Calvin Woon-Loong Chin^26^, Yan Yan Chen^14^, Boon Jun Chia^14^, Sonia Davila^22^, Jun Hui Goh^13^, Patricia Gomez^11^, Mar Gonzalez-Porta^6^, Pierre-Alexis Goy^6^, An An Hii^28^, Justin Jeyakani^6^, Saumya S, Raden Indah Kendarsari^10^, Raghavan Lavanya^14^, Jyn Ling Kuan^5^, Zihui Li^6^, Hengtong Li^15,16^, Jian Jun Liu^20,25^, Weng Khong Lim^22,23,28,29^, Mingxuan Lin^5^, Joseph Yio Linao^13^, Daniel Martana^10^, Vanessa Mok^4^, Ee Ling Ng^12^, Daryl Jun Xian Ong^13^, Qingsheng Peng^14,30^, Charumathi Sabanayagam^14,17^, Da Soh^14,16^, Helen Speirs^4^, Jin Su^13^, Zulkhairil Suradi^13^, Amelia Sutikna^6^, Wee Kiat Tan^13^, Joanna HJ Tan^6^, Yih Chung Tham^14,15,16,17^, Sahil Thakur^14^, Roberto Tirado-Magallanes^6^, Jonathan Tjendana^10^, Wilson Chandra Tjhi^10^, Cong Ling Teo^14^, Zhi Tien Yin Wong^14,31^, Yen Lynn Wong^10^, Can Can Xue^14^, Gretchen Weightman^4^, Chee Jian Pua^27^, Yee Yen Sia^25^, Tin Aung^14,17^

^25^Human Genetics, Genome Institute of Singapore (GIS), Agency for Science, Technology and Research (A*STAR), 60 Biopolis Street, #02-01 Genome, Singapore 138672, Republic of Singapore, ^26^Department of Cardiology, National Heart Centre Singapore, Singapore, 169609, Singapore, ^27^National Heart Research Institute Singapore, National Heart Centre Singapore, Singapore, 169609, Singapore, ^28^SingHealth Duke-NUS Genomic Medicine Centre, Singapore, 168582, Singapore, ^29^Laboratory of Genome Variation Analytics, Genome Institute of Singapore, Agency for Science, Technology and Research, Singapore, 138672, Singapore, ^30^Clinical and Translational Sciences Program, Duke-NUS Medical School, Singapore, ^31^Tsinghua Medicine, Tsinghua University, Beijing, China.

## Additional information

Supplementary Notes extensively describe and detail the infrastructure, operational, and human resource components of the PRECISE-SG100K Genomic Engine. We make a deep dive into the Stress Test, optimizations, and evaluate workflow recommendations aimed to improve overall processes.

