## Supplementary material for "Scaling Precision Medicine: Singapore’s PRECISE-SG100K Genomic Engine": Standalone Supplemental Notes referenced in Summary Manuscript

### Supplementary Notes

### Authors and affiliations

Claire Bellis,[1,2,3‡**] Gabriel Kolle,[4‡] Jacklyn Yong,[5‡] Maxime Hebrard,[6,3,7‡] Bitong Clarabelle Alexandrine Lin,[1] Tat Hung Koh,[1] Paul CP Cheng,[1] Shakita D/O Sunmugam,[1] Khai Koon Heng,[8] Zhicheng Xie,[8] Wee Yang Meah,[8] Xiao Yin Chen,[9] Erin Yi Ting Lim,[7] Jyn Ling Kuan,[5] Rudi Alberts,[5] Helen Speirs,[4] Sherilyn Lim,[5] Yen Lynn Wong,[10] Siew Hong Leong,[10] Jack Ling Ow,[6] Rodrigo Toro Jimenez,[6] Eleanor Wong,[11] Terry Yoke Yin Tong,[12] Swat Kim Kerk,[12] Jiali Yao,[13] Miao Ling Chee,[14] *PRECISE-SG100K Genomic Engine Consortium*, Khung Keong Yeo,[15,16] Ching-Yu Cheng,[14,17,18,19] Xueling Sim,[13] Weiling Zheng,[3] Chiea Chuen Khor,[1,8,9] Shih Wee Seow,[3] E Shyong Tai,[3,13,16,20] John C Chambers,[3,12] Nicolas Bertin,[6,3**] Patrick Tan[3,21,22,23,24*]

### Contact information

Correspondence and requests for materials should be addressed to:

**Dr. Claire Bellis EXTRACT GIS:

**Dr. Nicolas Bertin GRIDS:

‡Authors contributed equally to this work

### Affiliations

[1]EXTRACT Platform, Genome Institute of Singapore (GIS), Agency for Science, Technology and Research (A*STAR), 60 Biopolis Street, #02-01 Genome, Singapore 138672, Republic of Singapore, [2]Centre for Genomics and Personalised Health, Genomics Research Centre, School of Biomedical Sciences, Queensland University of Technology, Brisbane, Queensland, Australia, [3]Precision Health Research, Singapore, 139234, Singapore, [4]Illumina Australia and New Zealand, Level 3, 535 Elizabeth Street, Melbourne, Vic 3000, Australia, [5]Illumina Singapore, 11 Biopolis Way, #09-05 Helios, 138667, Singapore, [6]Genome Research Informatics & Data Science Platform, Genome Institute of Singapore (GIS), Agency for Science, Technology and Research (A*STAR), 60 Biopolis Street, #02-01 Genome, Singapore 138672, Republic of Singapore, [7]Garvan Institute of Medical Research, Sydney, New South Wales, Australia, [8]Integrated Genomics Platform, Genome Institute of Singapore (GIS), Agency for Science, Technology and Research (A*STAR), 60 Biopolis Street, #02-01 Genome, Singapore 138672, Republic of Singapore, [9]Laboratory of Complex Disease Genetics, Genome Institute of Singapore (GIS), Agency for Science, Technology and Research (A*STAR), 60 Biopolis Street, #02-01 Genome, Singapore 138672, Republic of

Singapore, [10]Illumina Singapore, 29 Woodlands Industrial Park E1, North Tech, Lobby 3, #02-13/18, 757716, Singapore, [11]NPM Programme Coordinating Office, Genome Institute of Singapore (GIS), Agency for Science, Technology and Research (A*STAR), 60 Biopolis Street, #02-01 Genome, Singapore 138672, Republic of Singapore, [12]Population and Global Health, Lee Kong Chian School of Medicine, Nanyang Technological University, 308232, Singapore, [13]Saw Swee Hock School of Public Health, National University of Singapore and National University Health System, Singapore, Singapore, [14]Singapore Eye Research Institute, Singapore, 169856, Singapore, *PRECISE-SG100K Genomic Engine Consortium*, [15]Department of Cardiology, National Heart Centre Singapore, Singapore, Singapore, [16]Duke-NUS Medical School, 169857, Singapore,[17]Centre for Innovation & Precision Eye Health, Yong Loo Lin School of Medicine, National University of Singapore, Singapore, [18]Department of Ophthalmology, Yong Loo Lin School of Medicine, National University of Singapore, Singapore, [19]Ophthalmology & Visual Sciences Academic Clinical Program (Eye ACP), Duke-NUS Medical School, Singapore 169856, Singapore, [20]Department of Medicine, Yong Loo Lin School of Medicine, National University of Singapore, 119228, Singapore, [21]Genome Institute of Singapore (GIS), Agency for Science, Technology and Research (A*STAR), 60 Biopolis Street, #02-01 Genome, Singapore 138672, Republic of Singapore, [22]SingHealth Duke-NUS Institute of Precision Medicine, 169609, Singapore, [23]Cancer & Stem Cell Biology Program, Duke-NUS Medical School, 169857, Singapore, [24]Cancer Science Institute of Singapore, National University of Singapore, 117599, Singapore.

### Supplementary Note 1: Singapore National Precision Medicine Program Phase I – Lessons Learned

Singapore's National Precision Medicine (NPM) program began with Phase I (SG10K_Health), which sequenced 10,000 Singaporeans to generate an Asian reference panel and establish early operational models, specifically in the area of investigating the prevalence on clinically actionable genetic variation. While scientifically successful, Phase I revealed multiple bottlenecks and rate-limiting steps that hindered scalability. To deliver Phase II at tenfold higher throughput, it was therefore essential to systematically capture these lessons and redesign operations. Key insights informed the development of the PRECISE-SG100K Genomic Engine, co-developed under a public-private partnership (PPP) between PRECISE, Illumina (ILMN), and a local sequencing service provider. Table 1 summarizes Phase I challenges, the learnings derived, and how these were incorporated into the design principles and infrastructure for Phase II.

*Table 1. Lessons learned from Singapore National Precision Medicine Program (NPM) Phase I (SG10K_Health). Challenges and risks encountered during pilot implementation are mapped to operational learnings and specific design features adopted in Phase II (PRECISE- SG100K Genomic Engine).*

| **Barriers to Upscaling of Genomics Capacity** | **Learning(s)** | **Phase II Recommendations and Set-up** |
|---|---|---|
| **DNA** | | |
| Various biospecimen types presenting as sample input for DNA extraction. | Standardization of biospecimen input type will enable automated DNA extraction at scale. | Acceptance of predominantly whole blood samples from multiple SG100K cohorts which were processed in an automated DNA processing facility<br>Supplementary Note 2: Singapore National Precision Medicine Program – Background<br>Figure 1 ***Box A*** |
| High variability in gDNA samples quality and quantity observed, due to the DNA extraction and sample normalization performed by the various participating cohorts using various combinations of assays and quantification equipment. | Centralization of DNA extraction at scale will reduce variability with one standard operating protocol. | A purpose-built, centralized and automated DNA processing facility, known as EXTRACT Lab, was established, with the capacity to generate 960 sequencing-ready samples weekly. Production-scale operations required minimally 768 samples/week.<br><br>Supplementary Note 3: EXTRACT Lab Module – Workflow, Biospecimens, and Quality Control<br>Figure 1 ***Box B*** |
| SG100K metadata acquisition through a manual process prone to user errors without a formal audit trail. | Automation will reduce user error and provide audit trail to allow for root cause identification. | Standardization of metadata format and ingress from SG100K via LabLink portal submission directly to Illumina Clarity LIMS (***CLIMS***).<br><br>CLIMS which tracks samples and workflow processes was deployed at both DNA module and IGA.<br>Supplementary Note 4. Illumina Data Generation Infrastructure<br>Figure 1 Illumina Data Generation Infrastructure |
| Absence of laboratory information management system (LIMS). | A LIMS that allows sample and workflow tracking will be needed in scaled up operations. | |
| Error-prone manual library preparation and sequencing. | Automated library preparation and sequencing will enable higher throughput with fewer user errors. | Deployment of the Illumina Data Generation Infrastructure (IDGI), an automated library preparation and sequencing workflow.<br>Checkpoints which serve as quality control steps were deployed to prevent cascade of potential issues in the event of workflow deviation.<br>Supplementary Note 4. Illumina Data Generation Infrastructure |

| Barriers to Upscaling of Genomics Capacity | Learning(s) | Phase II Recommendations and Set-up |
|---|---|---|
| | | Figure 1 ***BOXES C_E*** |
| Challenges in troubleshooting with deployment of manual workflows. | Manual touchpoints had to be reduced via workflow automation.<br>Workflow checkpoints will be helpful in troubleshooting and triaging of issue.<br>Process for managing sample and operational issues needs to be well established. | Work processes and data management processes have been automated as much as possible to reduce manual interventions and enhance operational efficiency.<br>Workflow checkpoints were established with the aim of triaging samples that do not pass criteria.<br>Supplementary Note 6: Operational Considerations in the PRECISE-SG100K Genomic Engine |
| **Data** | | |
| Data provenance and governance policies. | | Sequencing data is managed with Illumina Connected Analytics (***ICA***), a secured platform for data storage, management, and analysis.<br>Quality control steps based on specific data metrics served as checkpoints to stop the automated data analysis process and trigger a failure notification and initiate a review.<br>Supplementary Note 4. Illumina Data Generation Infrastructure<br>Figure 1 ***BOXES C-E*** |
| Manual sample tracking process prone to user errors. | Automation of the process will minimize workflow deviations and reduce cascade effects of such deviations | Establishment of PRECISE Sample Tracking System (***STS***) which generated de-identified unique sample identifier (NPMID) carried over by all parties of the Genomic Engine.<br>Supplementary Note 5. Genome Research Informatics & Data Science - Informatics and data stewardship<br>Figure 1 ***BOXES F and G*** |
| Challenges in increasing data production in terms of storage availability on premise. Hardware maintenance and data redundancy measures are costly. | Storage on cloud is virtually unlimited. Cloud provider guarantees availability and zero data loss. | Sequencing data generated are streamed to cloud (Figure 1 **BOX C**) with all downstream analysis maintained and archived on cloud (Figure 1 **BOXES E, H, and I**)<br>Supplementary Note 4. Illumina Data Generation Infrastructure<br>Supplementary Note 5. Genome Research Informatics & Data Science - Informatics and data stewardship |
| Evolving legislation on genomic data sensitivity impose additional measures in term of security and governance | Cloud provider maintains state-of-the-art high security infrastructure | |

| Barriers to Upscaling of Genomics Capacity | Learning(s) | Phase II Recommendations and Set-up |
|---|---|---|
| **Overall infrastructure** | | |
| Uncertainty in accurate and timely dissemination of information to workgroups. | Regular meeting cadence for workgroups will permit forward planning, securing timeslots for key stakeholders and their representatives. Standardizing meeting agenda and documenting key decisions in a centralized and accessible location for all will ensure transparency and avoid information asymmetry. | Regular meeting cadence with various workgroups and Microsoft Teams as main communications channel was established with distribution and alignment of meeting minutes.<br><br>Supplementary Note 6: Operational Considerations in the PRECISE-SG100K Genomic Engine |
| Ad-hoc assembly of operational team members to investigate and diagnose deviations related to genome quality metrics. | A mechanism to capture and document decisions related to samples which deviated from the happy path with visibility to key stakeholders was required. | The implementation of a formal ***Sample Review Clinic*** meeting every 2 weeks provided the opportunity to discuss and resolve problematic samples, monitor trends, and inform of any deviations occurring which may impact production.<br><br>Supplementary Note 6: Operational Considerations in the PRECISE-SG100K Genomic Engine |
| Absence of established system for operation/trends monitoring. | An established system of data trends monitoring, and rapid post-sequencing QC based on primary and secondary analysis will serve as an early alert system on existing or potential operational issues, allowing workgroups with sufficient lead time to work on mitigation plans to avoid or minimize impact to production. | A SG100K dashboard was built to enable day-to-day review of the ongoing operation and the use of DRAGEN in ICA on AWS cloud allowed for fast and high throughput (240 samples in 13-15 hours) turnaround, both of which enabled quick review of trending data metrics. Any trends that were unexpected would be flagged and communicated to the workgroups via the communication channel established.<br><br>Supplementary Note 5. Genome Research Informatics & Data Science - Informatics and data stewardship<br><br>Figure 1 **BOX G** |
| Phase I held quarterly meetings with key stakeholders providing updates on deliverables. | | Phase II required a structured ecosystem-wide mechanism for oversight. The ***Joint Steering Committee*** (JSC) was convened every six months to review project goals, timelines, and progress. In the case |

| Barriers to Upscaling of Genomics Capacity | Learning(s) | Phase II Recommendations and Set-up |
|---|---|---|
| | | of situations which were not solved at the SRC level, these would be raised for discussion and resolution at JSC.<br><br>Supplementary Note 6: Operational Considerations in the PRECISE-SG100K Genomic Engine |

**Supplementary Note 2: Singapore National Precision Medicine Program – Background**

Precision Medicine (PM) is a data-driven approach to healthcare focusing on understanding how individual and group genetic variation can interact with environmental and lifestyle factors to influence health and disease. When appropriately applied, PM has the potential to deliver more accurate and effective disease treatment and prevention strategies, mitigate rising healthcare costs while maintaining good clinical outcomes, and catalyze a wide range of economic activities in the genomics and biomedical sectors. To reap PM's benefits, many governments globally have initiated national PM programs such as All of Us[1, 2] (formally Precision Medicine Initiative[3]), Genomics England[4-6], UK BioBank[7-12], Our Future Health[13], Estonia Biobank[14], Taiwan Biobank[15] and Precision Medicine Initiative[16], DeCODE[17], FinnGen[18], China Kadoorie Biobank[19], BioBank Japan[20], Tohoku Medical Megabank Project[21], Pan-Canadian Genome Library[22, 23], Hong Kong Genome Project[24], Australian Genomics[25, 26], Emirati Genome Project[27] Egypt Genome Project[28] and Qatar Genome Program[29-31]. Several PM programs are also being initiated at the regional and health system level, including H3Africa Consoritum[32],eMERGE[33], DiscovEHR/MyCode[34-37], Healthy Nevada Project[38, 39], Bio*ME*[40], BioVU[41-43], OneDukeGen[44], and Global BioBank Meta[45]. Core to each of these programs is the mandate to translate genomic findings into clinically relevant PM interventions[46-48]. Complementing these programs are global initiatives promoting data harmonization and exchange such as the Global Alliance for Genomics Health (GA4GH)[49], International Health Cohorts Consortium (IHCC)[50] and Global Genomic Medicine Consortium (G2MC)[51]. Such international programs are essential to address serious inequities in global PM research, as the majority of the world's PM data is focused on Euro-centric population[25, 52-60]. On a technical level, one of the key factors to the success of such initiatives is a robust genomics data generation platform that balances financial cost, resource utilization (including biospecimen use), data quality, and sample throughput.

Singapore, with a total population size of approximately 5.92M and a resident population size of 4.15M[61], has one of the highest population densities (at >7,500 residents per square km), globally[62]. The Singapore population has a rich and heterogenous multi-ancestry composition of three major groups: Chinese (74.2%), Malay (13.7%), and Indian (8.9%) sculpted by regional migration and genomic admixture[63, 64]. Singaporeans are living longer - one third of Singaporean's population will be over 65 by 2050[62], and in 2020-2025 the average Singaporean life expectancy is approximately 13% higher than the global average (84.1 v's 73.3)[65, 66]. In 2017, Singapore launched its National Precision Medicine (NPM) program[63], aiming to collect and integrate genomic, phenotypic, and clinical data for up to 10% of its resident population. Coordinated by PRECISE (Precision Health Research, Singapore), NPM is structured as a three-phase initiative. NPM Phase I assembled an Asian reference genome cohort of 10,323 healthy consented individuals (SG10K_Health[63],[64]). While successfully completed, Phase I was performed using pre-existing heterogenous infrastructure and labour-intensive lab-scale workflows (Supplementary Note 0). In planning for Phase II, 100,000 whole-genome sequence (WGS), an after-action review of Phase I highlighted four key areas for efficiency improvements to be addressed to achieve the desired ten-fold scale up in NPM Phase II:

1) High variability in gDNA sample quality and quantity necessitated repetitive manual interventions to standardize samples. This was due to DNA extractions being performed by the various participating cohorts.
2) Metadata acquisition and sample tracking processes was performed via email and manual spreadsheet capture which was unscalable particularly in the absence of an advanced laboratory information management system (LIMS).
3) Manual library preparation and sequencing operations introduced various errors requiring significant root-cause investigations and process deviations.
4) Data storage and analysis was performed on-premises where limitations involving computation efficiency, hardware maintenance and storage requirements were identified.

Here, we share an implementation blueprint and the preliminary results for the 102,121 WGS of Singapore's NPM Phase II PRECISE-SG100K Genomic Engine, detailing key features, operational parameters, and critical workflow aspects. We believe the insights from this report describing the Singapore experience will provide highly valuable operational insights to nascent PM initiatives globally.

**PRECISE-SG100K Population Cohort Biospecimens**

The SG100K population study is a longitudinal project with on-going participant enrolment comprised of four cohorts: Healthy for Life in Singapore (HELIOS) at Lee Kong Chian School of Medicine, Nanyang Technological University[67]; Singapore Population Health Study (SPHS) at Saw Swee Hock School of Public Health, National University of Singapore; Singapore Epidemiology of Eye Diseases study (SEED) at Singapore Eye Research Institute (SERI), SingHealth; and SingHeart study at National Heart Centre Singapore, SingHealth.

To enter the PRECISE-SG100K Genomic Engine, all samples from consented SG100K participants were deidentified with sharing metadata restricted to the participants' self-reported sex which was compared to the sequence-inferred sex ploidy as an initial QC checkpoint during downstream sequencing analyses. Approximately 90% of biospecimens were stored as whole blood. The remainder were considered as exotic samples since their processing required bespoke, non-automated DNA processing operations. Examples of exotic samples included previously extracted (legacy) DNA, buffy coats, and blood clots. Contributions from HELIOS and SPHS constituted most of the samples (>90%). By restricting the majority of incoming biospecimens to whole blood in Phase II, it became possible to achieve the necessary throughput to sequence 100K genomes within the 3-year program timeline, at an average rate of approximately 800 DNA samples weekly.

Logistical coordination of SG100K sample acquisition was centrally undertaken by EXTRACT. Each SG100K sample site was independently engaged by EXTRACT to onboard team members for biospecimen sample and metadata submission. EXTRACT supplied the SG100K sites with compatible consumables (96-deep well pre-barcoded plates and seals) for standardized biospecimen transfer protocols. To reduce variability in the whole blood sample input, team members at SG100K sample sites were provided guidance on the preparation of biospecimens prior to submission to EXTRACT.

### PRECISE-SG100K Population Cohort Demographics

The SG100K population reflects Singapore's multi-ethnic composition, with participants enrolled from HELIOS, SPHS, SEED, and SingHeart cohorts. Across the 102,121 genomes sequenced, the ethnic distribution broadly mirrors national population proportions: ~74% Chinese, ~14% Malay, ~9% Indian, and ~3% other ancestries. Sex distribution was balanced, with ~50% male participants across cohorts. Age at baseline varied by cohort and ethnicity (Table 2), with mean values spanning mid-40s to early-60s, thereby capturing both early- and late-adult life stages relevant to disease onset and progression. This diverse representation provides a powerful foundation for investigating ancestry-specific disease risk and gene-environment interactions in Asian populations.

| **Cohort** | **Chinese** | | **Indians** | | **Malays** | | **Male %** | | **Totals** | |
|---|---|---|---|---|---|---|---|---|---|---|
| | *n* | Age (± SD) | *n* | Age (± SD) | *n* | Age (± SD) | *n* | Age (± SD) | *n* | Age (± SD) |
| Overall | 75,373 | 52.0 (13.3) | 16,250 | 49.5 (13.6) | 13,545 | 46.8 (14.1) | 45,527 | 51.2 (14.1) | 107,139 | 51.1 (13.6) |

***Table 2. Demographic characteristics of the PRECISE-SG100K population cohort, stratified ethnicity.*** *Baseline characteristics include sample size (n), distribution of ethnic groups (Chinese, Indian, Malay, Other), sex distribution (% male), and mean age at baseline with standard deviation (years). Overall summary values reflect the combined cohorts.*

### Overview of PRECISE-SG100K Genomic Engine

The Genomic Engine was co-developed via a public-private partnership between PRECISE, the Genome Institute of Singapore (GIS), Illumina, and a local sequencing provider. This collaboration integrated public sector program management and cohort engagement with commercial capabilities in high-throughput sequencing, automation engineering, and informatics. The platform’s modular architecture (Figure 1, Main Text) comprises:

1. DNA Extraction – EXTRACT Module: Centralised high-throughput laboratory for standardised processing of predominantly whole-blood samples, ensuring consistent yield and purity.
2. Automated Library Preparation & Sequencing – Illumina Data Generation Infrastructure (IDGI): Robotics-enabled workflows linked to NovaSeq instruments, supporting sustained weekly throughput.
3. Cloud-based Informatics & Data Management – Genome Research Informatics & Data Science (GRIDS): Secure, standards-aligned data processing, and quality control, with integration into the national TRUST platform.
4. Sample Tracking System (STS): AWS-hosted service linking de-identified unique sample IDs to metadata, QC metrics, and operational status across all modules.

An “automation-first” design reduced manual touchpoints, accommodated upstream biospecimen variability, and ensured downstream readiness for both research-grade and future clinical-grade pipelines. All modules were integrated via Illumina Clarity LIMS (v 6.2.1.11) and STS for real-time process monitoring and full auditability.

### Supplementary Note 3: EXTRACT Lab Module – Workflow, Biospecimens, and Quality Control

#### Summary of Performance to Metrics

The core intake and DNA extraction module, termed EXTRACT, was established as a high-throughput, automation-first facility to standardise processing of biospecimens for the PRECISE-SG100K project. Over the course of the programme, 102,469 biospecimens were provisioned from cohort sites and processed through EXTRACT, encompassing whole blood, legacy DNA, clotted blood, and buffy coat inputs. DNA was successfully extracted, normalized, and qualified for sequencing for most submissions, with pass rates exceeding predefined thresholds across input types. EXTRACT achieved a steady-state throughput of ~960 samples per week, with the bulk of processing occurring during 2023-2024 when sample accrual was at maximum scale. Governance via predefined QC metrics and continuous process optimization enabled consistent delivery of sequencing-ready DNA, ensuring compatibility with downstream library preparation and sequencing pipelines.

#### Biospecimen Reception Format

Cohort sites were supplied with pre-barcoded 96-well plates, each well aliquoted with 500µl of whole blood, accompanied by digital manifests uploaded via LabLink, a secure web-based submission interface for metadata. Plates were visually inspected on site before transfer to the EXTRACT team, with chain of custody maintained for rectification in the event of anomalies (e.g., missing samples or compromised seals). Upon receipt, plate identifiers and sample-level metadata were automatically ingested into Illumina Clarity LIMS (CLIMS), where each sample was assigned a de-identified persistent identifier (NPMID) carried through all downstream processes.

**Workflow Design and Capacity**

Automated extraction of SG100K whole blood samples was performed on Explorer G3 workstations using the Revvity chemagic™ Prime DNA Blood 400 Kit H96, generating 96 sequencing-ready DNA samples in ~2.5 hours of run time (excluding thawing, reagent preparation, and maintenance). Assays were programmed through plate::works™ automation scheduling software, with customizations introduced to improve yield and quality: (i) a starting input of 500µl whole blood per sample; (ii) division of each sample into two 250µl aliquots for lysis; (iii) adjustment of lysis reagent volume to 300µl per 250µl aliquot (vs. 500µl for 500µl input in the standard protocol); and (iv) extension of the lysis incubation step by 100 seconds, to a total of 400 seconds. Daily processing of two 96 sample plates established an initial steady-state throughput of ~960 sequencing-ready samples per week, sufficient to meet project timeline target.

DNA quality control (QC) was implemented as a central pillar of the PRECISE-SG100K Genomic Engine, ensuring that biospecimens processed by the EXTRACT Module were consistently suitable for downstream sequencing (Supplementary Note 4. Illumina Data Generation Infrastructure). Quantification was performed on the extraction eluate to determine DNA concentration, with the values automatically fed into an internal algorithm that calculated the precise DNA and buffer volumes required to normalize all samples to a target concentration, volume, and total DNA amount (detailed below).

Following normalization, an exit QC was performed to confirm that targets had been achieved, and results were shared with the downstream sequencing provider via the LabLink portal in CLIMS to inform library preparation considerations. All QC and normalization metadata were synchronized with the Sample Tracking System (STS; Supplementary Note 5. Genome Research Informatics & Data Science - Informatics and data stewardship), providing a live audit trail across the Genomic Engine.

**Primary QC parameters**

The QC workflow integrated automated laboratory measurements with informatics systems for traceability, auditability, and operational oversight. DNA eluates were quantified using fluorescence-based (Qubit, VICTOR Nivo™) assays and assessed for purity by absorbance spectroscopy (Unchained Labs, Lunatic). The passing criteria were defined as:

- Amount ≥1.5µg DNA
- Volume ≥45µl
- Concentration ≥20ng/µl
- OD 260:280 between 1.8-2.0
- OD 260:230 between 2.0-2.2

The operational targets were set higher; 2.4µg in 61µl at 40ng/µl, to provide sufficient DNA template redundancy for the generation of two independent library preparations.

93.6% of DNA samples qualified as PASS when the 260:230 absorbance ratio was excluded from the criteria (Table 3 and Figure 2 Box E). Based on the distribution of A260/230 that was generated, we do not see an impact of the ratio on the generation of high quality WGS using the Genomic Engine sequencing protocols.

***Table 3. Quality Control Metrics and Pass Rates***

| Quality Control Metric | Pass Criteria | Percent Passed (%) |
|---|---|---|
| **DNA concentration** | >20ng/µl | 98.6 |
| **A260/280** | 1.8-2.0 | 96.2 |
| **A230/260** | 2.0-2.2 | 42.5 |
| **DNA volume** | >45µl | 99.8 |
| **DNA amount** | 1.5µg | 97.7 |

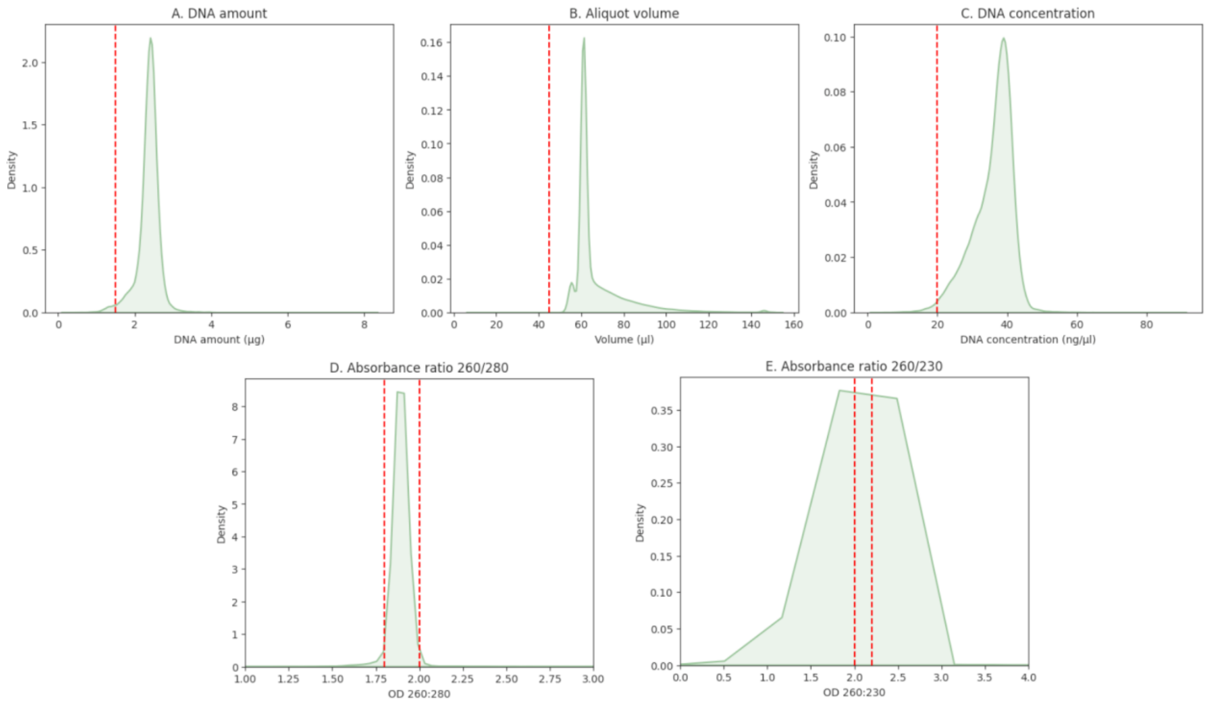

***Figure 1. Distribution of DNA quality metrics for 102,469 samples.*** *A-E. Density plots for A) DNA amount, B) Aliquot volume, C) DNA concentration, D) Absorbance ratio 260/280, E) Absorbance ratio 260/230. Quality thresholds are show as red dotted lines.*

### Biospecimen Input Type

While the majority of SG100K samples were freshly collected whole blood (Figure 3), the PRECISE-SG100K Genomic Engine was intentionally designed to accommodate non-standard or "*exotic*" input types, reflecting the heterogeneity of real-world cohorts and legacy collections. These included archived DNA, clotted blood, and historically frozen biospecimens, each presenting unique technical challenges to extraction and downstream sequencing.

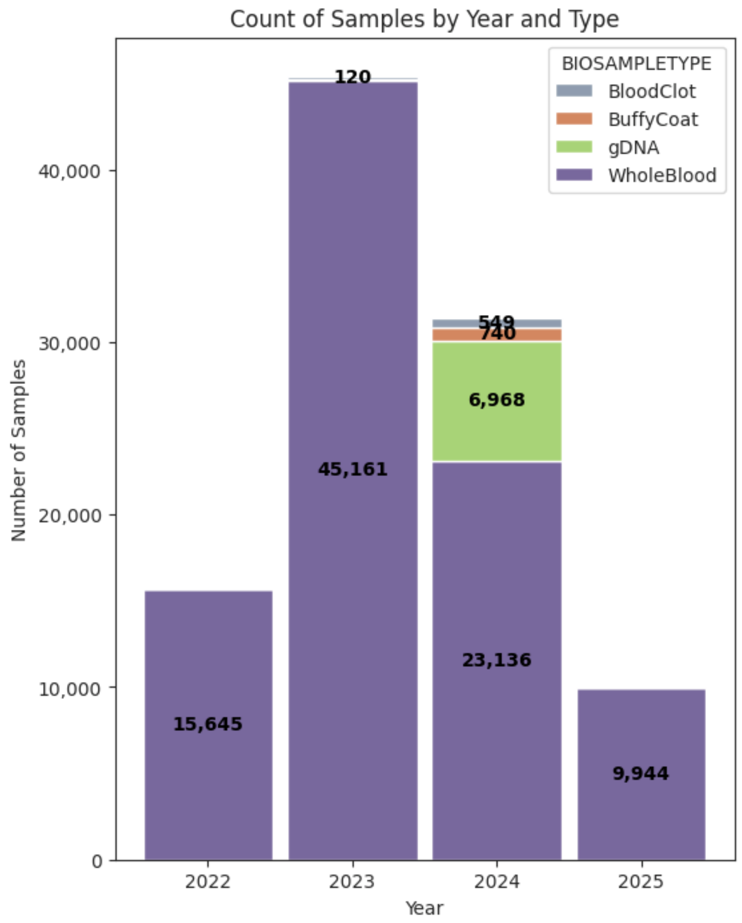

***Figure 2. Breakdown of sample provisioning from SG100K cohorts to EXTRACT Lab over project lifecycle.*** *Individual counts are not displayed when a category has < 100 samples. In 2023, 95 gDNA were provided. In 2025, 16 gDNA and 95 Buffy Coat were provided.*

Over 7,000 previously extracted DNA samples were submitted to the Genomic Engine. QC thresholds for archived DNA followed the same criteria as freshly extracted samples with preparation of DNA submission plates run on a modified version of the whole blood QC-normalization-QC workflow (Figure 4).

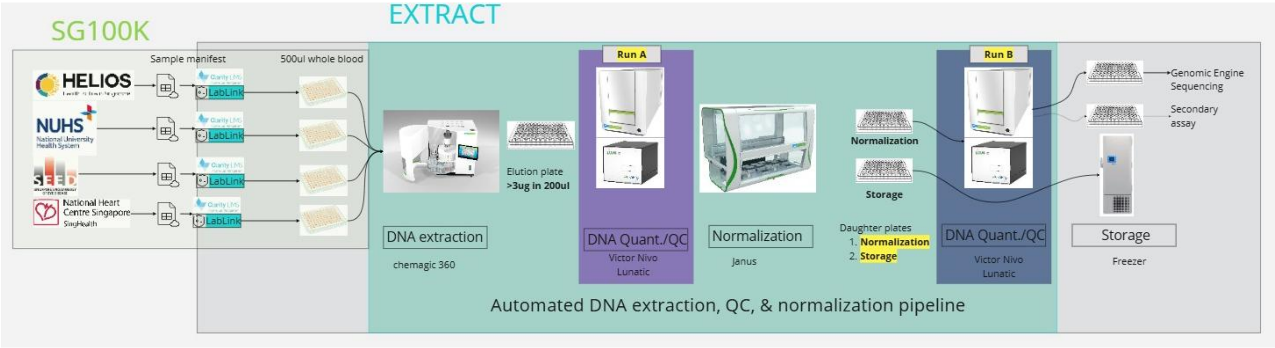

***Figure 3. EXTRACT Lab whole blood operational workflow.*** *Over 90% of SG100K samples were provided to EXTRACT Lab as whole blood biospecimens. Operationally, this approach allowed for scale-up of operations due to standardization of the input material, a learning lesson from NPM Phase I. Two DNA plates were generated for each 96-deep well whole blood plate; 1) sequencing-ready normalized DNA submitted to sequencing provider and 2) any excess DNA template was stored in a stock DNA plate.*

Approximately 700 submissions originated from a cohort in which a subset of samples were intentionally collected as clotted blood under the study protocol. These were processed in the EXTRACT Lab through a customised, predominantly manual lysis protocol developed to maximize quality DNA recovery. While QC outcomes showed slightly lower yields than freshly collected whole blood, most clotted samples still met the requirements for downstream sequencing, with a 100% success rate for sequencing quality metrics.

Like the clotted blood cohort, another cohort provided ~850 samples as buffy coat derived from whole blood. Given the high white blood cell content, iterative optimization assays were required to establish the optimal input volume, of 250µl, without overloading the magnetic beads. Once stabilization protocols were agreed upstream with the cohort to preserve remaining material for other assays, buffy coat samples were successfully processed through a modified automated extraction workflow, achieving a 100% success rate for WGS data bundle delivery.

Stabilization protocol: addition of 1,000µl Rnalater to 500µl of buffy coat, mix before aliquotting 250ul into pre-barcoded 96 deep-well plate.

**Intentional Test Plate**

Because cohort samples were not processed sequentially and biospecimen input types were introduced at different times, downstream operations incorporated deliberate test plate runs. These were monitored in real time by all stakeholders to validate workflows before bulk cohort samples were prepared and submitted. Importantly, this approach provided assurance of process robustness without adversely affecting turnaround times.

**Scalability and governance**

By incorporating flexible protocols and maintaining detailed metadata through CLIMS and STS, the Genomic Engine ensured that non-standard samples were tracked, reviewed, and adjudicated with the same rigor as standard inputs. In the event where WGS QC metrics deviated, Sample Review Clinics (SRCs) provided a structured forum, for stakeholder consensus on whether to reprocess, resequence, or exclude (abandon) the sample (Supplementary Note 6: Operational Considerations in the PRECISE-SG100K Genomic Engine).

Overall, the ability to incorporate archived DNA, clotted blood, and buffy coat samples expanded the Genomic Engine's applicability beyond freshly collected whole blood, increased resilience to upstream variability, and established a framework for future Phases where sample diversity may broaden further.

**Key Considerations for Automated High-throughput DNA Extraction**

1. **Define and enforce QC metrics.**
   Establishing standardized DNA QC PASS thresholds (≥1.5μg in ≥45μl, ≥20ng/μl, OD 260:280 of 1.8-2.0, OD 260:230 of 2.0-2.2) ensured biospecimens processed through EXTRACT were consistently suitable for sequencing across **n = 102,469 samples**.

2. **Automate with targeted optimization.**
   Automated workflows delivered steady throughput (~960 samples/week). Protocol modifications, including split aliquots and extended lysis, improved yields from frozen and variant inputs.

3. **Implement structured governance.**
   Real-time monitored test plates validated workflows for new cohorts or input types, while bi-weekly Sample Review Clinics provided a forum for deciding re-extraction, resequencing, or exclusion of QC outliers. This framework enabled successful processing of diverse inputs; including >7,000 archived DNA samples, <900 clotted blood samples, and buffy coat submissions, all meeting sequencing requirements.

### Supplementary Note 4. Illumina Data Generation Infrastructure

#### Summary of Performance to Metrics

The core module for genomic data generation termed the Illumina Data Generation Infrastructure (IDGI), was built as a highly scalable system with a goal of delivering over 100,000 whole genome sequences (WGS) within the project timeframe of 3.25 years. Overall, the IDGI received 102,618 DNA samples in 1,094 96-well plates and delivered sequencing results for 102,202 samples (99.59%). The median time from submission of extracted DNA to successful transfer of data was 11 days. A maximum of 39,927 samples were delivered in a single calendar year with the majority of samples processed in 2023 and 2024, where sample submission and processing were at maximum capacity and throughput. This robust infrastructure, governed by quality thresholds and an automation-first principle, delivered 98.9% of samples on first pass.

##### Quality Control Metrics

For the SG100K, quality control (QC) metrics for successful data generation were agreed to ensure high quality WGS data delivery and to reduce the likelihood of suboptimal data that may confound the downstream use of the data (Table 4). These metrics were designed based on experiences from previous programs including the Genomics England 100,000 Genomes Project. Along with QC metrics that were established for incoming DNA samples from EXTRACT (Supplementary Note 3: EXTRACT Lab Module – Workflow, Biospecimens, and Quality Control), the IDGI established key QC criteria for sequencing data to ensure data integrity, including NovaSeq™ 6000 run QC to account for sequencer performance, post demultiplexing and FASTQ generation QC to account for sequencing yield per sample (Gbp), sex concordance to determine potential sample swaps and post alignment QC using a set of quality criteria defining sufficient coverage and read quality to ensure high quality variant calling.

*Table 4. Quality Control Metrics*

| Quality Control Metric | Process | Metric | Description |
|---|---|---|---|
| **DNA concentration** | Sample submission | >20ng/µl | The concentration of DNA as detected by fluorescence intensity (Qubit or PicoGreen). |
| **A260/280 ratio** | Sample Submission | 1.8-2.0 | Purity of DNA detected by UV absorbance. |
| **A230/260 ratio** | Sample Submission | 2.0-2.2 | Purity of DNA detected by UV absorbance. |
| **DNA volume** | Sample Submission | >45µl | The minimum volume required for high-throughput sequencing operations. |
| **DNA amount** | Sample Submission | 1.5µg | The amount of DNA submitted for sequencing (2.0µg during Stress Test). |
| **Instrument Run (flowcell) Yield** | Run Upload to BSSH | 2.4Tbp | NovaSeq 6000 flow cell yield lower threshold. Run yield below this results in resequencing of flow cell. |
| **Instrument Run (flowcell) Q30** | Run Upload to BSSH | 85% | NovaSeq 6000 flow cell number of bases >Q30. Triggers investigation. |
| **Flowcell sex mismatch** | BclConvert | 3 | Number of samples in a flow cell that do not match submitted sex. Triggers investigation. |
| **Raw Sample Yield** | BclConvert | 90Gbp | Number of raw bases for an individual sample. Triggers additional sequencing. |
| **Q30 bases > 85 excluding duplicates and trimmed bases** | DRAGEN analysis | 77.5Gbp | Number of bases that have a quality score above Q30 after excluding duplicate read pairs and trimmed bases. Triggers additional sequencing. |
| **Autosomal coverage > 15X** | DRAGEN analysis | 95% | Number of bases of the autosomal (chromosome 1-22) genome that has a coverage higher than 15X after read alignment. Triggers review. |
| **Autosomal callability** | DRAGEN analysis | 95% | Number of bases in the autosome that can be confidently called, based on DRAGEN variant caller assessment. Triggers review. |
| **Estimated contamination** | DRAGEN analysis | 1% | Estimated cross-individual contamination of the sample by another human source. Triggers review. |
| **Ploidy/sex match** | DRAGEN analysis | True | Comparison of DRAGEN estimated ploidy versus supplied sex (male XX, female XY). Any mismatch including sex aneuploidy triggers review. |

**IGDI Performance to Data QC Metrics**

Quality of sequencing data was assessed through yield measured as the total bases sequenced for a sample and the number of bases of Q30 excluding duplicates and trimmed reads; coverage of autosome bases at 15x, and the percentage of autosomal callability as a proxy for bases that could provide an accurate SNV call. Figure 5 shows the distribution of QC metrics on delivered samples in relation to the QC metric. Of the delivered samples, only 0.12% did not meet all of these metrics.

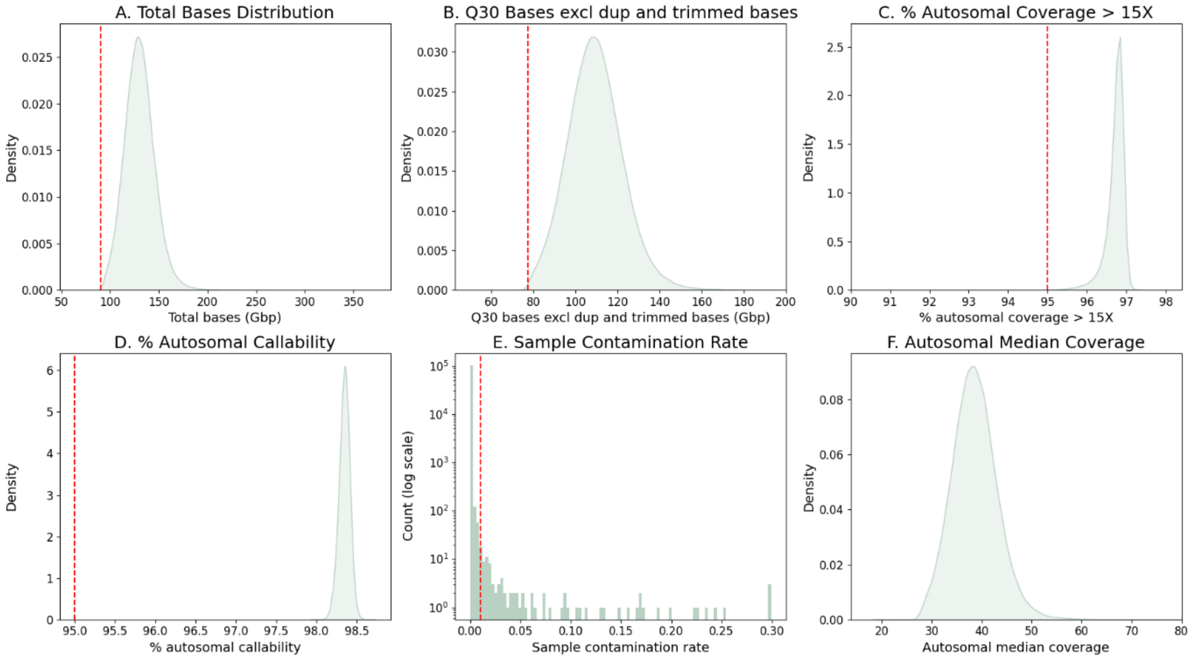

***Figure 4. Distribution of key quality control metrics for 102,202 samples**. A-D. Density plots for A) Total Bases (raw sequencing yield), B) Q30 Bases excluding duplicates and trimmed bases, C) Percentage autosomal coverage greater than 15 times, D) Percentage autosomal callability. E) Histogram of sample contamination rate, y-axis is on a log10 scale. F) distribution of sample coverage based on median coverage in autosomal regions. Quality thresholds are show as red lines. All quality metrics were output by DRAGEN (version 3.7.8).*

A key part of the IGDI setup was a focus on first pass efficiency to limit the number of samples that required additional sequencing, thereby ensuring cost efficiency. To that end, 98.9% of samples had sufficient yield and coverage QC metrics after a single sequencing round, meaning that those samples did not need further data generation. Of the remaining samples, we were able to add additional sequencing to reach QC metric thresholds (0.89% resequencing of the sample library, 0.15% re-library preparation and sequencing).

Another key metric we monitored was “cross sample contamination” by another human sample. Cross contamination can confound downstream analytics. We observed contamination for 0.09% of samples (Figure 5 Box E) of ≥ 1%. Of these, 58.9% occurred as a single instance in a plate of 96 samples, suggesting isolated contamination events, with 4 occurrences of contamination of 4 or more samples in a plate (maximum of 8 per plate).

We also put in place a system to identify potential plate level sample swap. This information was used to determine if there were any adverse events such as plate swaps, rotations, or incorrect metadata provision for samples. To do this, we compared the reported sex of the individual against the X and Y ploidy based on genomic data (see IDGI infrastructure and methodology). A threshold of three or more samples with a mismatch between observed ploidy (based on read mapping to the X and Y chromosomes) and expected ploidy for any 24 samples (the number of samples in a NovaSeq 6000 flow cell) was used to assess adverse events. Six plates were observed with a ploidy mismatch caused by incorrect incoming metadata and two plates with ploidy mismatch caused by a plate rotation. The ploidy mismatches enabled us to rapidly identify these adverse events and ensured that we could continue the process and deliver the samples.

Sex chromosome ploidy was also assessed as part of the individual sample QC process. 0.4% of samples were discrepant based on observed ploidy vs. expected ploidy (Table 5). 0.19% of samples had an unusual ploidy of X0, XXX, XXY, XXXY, or XYY. We observed that the X0 samples were most often male (86.8%) and were identified to have less than 50% of expected Y chromosome coverage. This is consistent with observations from other projects and likely caused by mosaic loss of Y chromosome in blood samples[1].

***Table 5. Sex chromosome ploidy for 102,202 samples***

| Ploidy | No. of samples | Percentage male (based on submitted metadata) | Percentage female (based on submitted metadata) |
|---|---|---|---|
| XY | 43,327 | 99.8 | 0.2 |
| XX | 58,684 | 0.2 | 99.8 |
| X0 | 114 | 86.8 | 13.2 |
| XXX | 21 | 0.0 | 100.0 |
| XXXY | 1 | 100 | 0.0 |
| XXY | 37 | 100 | 0.0 |
| XYY | 18 | 100 | 0.0 |
| total | 102,202 | 42.6 | 57.4 |

#### Turnaround Time

The IDGI was setup to ensure efficient, scalable and continuous data delivery (Figure 6). To that end, turnaround time (TAT), defined as the time taken between receipt of sample DNA plates to delivery of WGS data, was an important metric to ensure continuous data generation and delivery. The program far exceeded the TAT metrics set out at the commencement of the program, with 75% of samples delivered within 19 days, and 90% of samples delivered within 28 days. The shortest TAT was 54 hours, while the median was 11.19 days and 95% of samples were delivered within 42 days (Table 6).

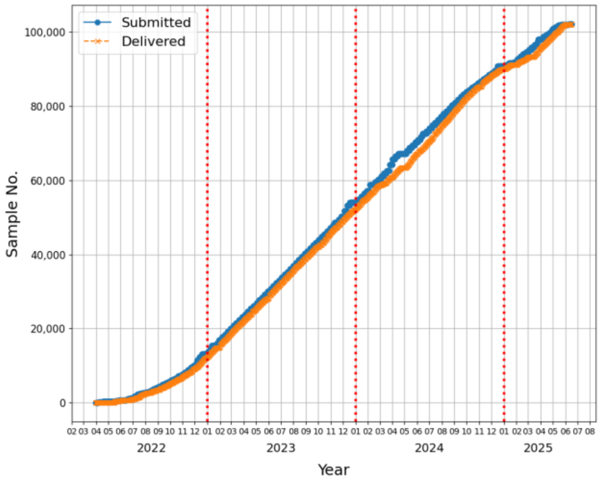

***Figure 5. Cumulative plot showing timing of submitted and delivered samples for SG100K project.*** *Samples ordered from 1-102,202 submitted as DNA plates (blue) and delivered as genomic data (orange) across the duration of the project (April 2022-June 2025).*

***Table 6. Final Metrics for Turnaround Time***

| Statistic | IDGI value | Target |
|---|---|---|
| Minimum | 2.33 days | NA |
| Median | 11.2 days | NA |
| 75th Percentile | 18.7 days | NA |
| 95th Percentile | 41.6 days | NA |
| % within 38 days | 93% | 75% |
| % within 60 days | 98% | 90% |
| Total samples | 102,202 | 100,000 |

**Key considerations for WGS data generation at scale**

Based on this project, we identified several key considerations to enable high volume WGS sequencing project:

1. Well defined QC metrics for input DNA and delivered data.
2. Process automation at all steps, with no manual intervention for samples passing QC metrics.
3. Rapid identification of failed samples or process failures and defined standard operating procedures (SOP) to handle these.
4. A sample review framework, here termed Sample Review Clinic (SRC), to discuss and confirm action on samples failing QC metrics that do not have clearly defined next steps under the SOP.

**IDGI Infrastructure and Methodology**

The Illumina Data Generation Infrastructure (IDGI) was designed to support high-throughput whole genome sequencing (WGS) with a modular and scalable architecture. It integrates automated wet lab workflows, cloud-based data analytics, and comprehensive metadata tracking to ensure reproducibility, traceability, and operational efficiency. The system spans the entire lifecycle of a sample, from DNA extraction at the EXTRACT facility to data delivery to the GRIDS data stewardship team, minimizing manual intervention and maximizing throughput.

The infrastructure comprised three components i) Illumina Genomics Architecture (IGA) for wet lab operations ii) the Illumina Data Analysis Module for secondary analysis and QC of data and iii) the Illumina Sample and Data Management Dashboard for project and data management (refer to Figure 1 in the main manuscript).

**i) Illumina Genomics Architecture**

Wet lab operations were governed by the Illumina Genomics Architecture (IGA), an automation-compatible workflow that orchestrates library preparation and sequencing. Sample manifests, containing de-identified IDs, the sex (male/female) of the individual and QC metrics, were submitted via Illumina Clarity™ LIMS (version 6.2.1.11). These initiate a controlled handover process within the EXTRACT facility to the local sequencing service provider, ensuring sample integrity through visual inspection prior to sequencing. Library preparation and pooling followed the Illumina DNA PCR-Free (IDPF) protocol, with samples processed in 96-well plates using the Hamilton MicroLab Star liquid handling system and sequenced on NovaSeq 6000 instruments. The IGA workflow utilized Clarity LIMS™ for sample tracking, interfacing with the Hamilton MicroLab Star liquid handling system and to govern the steps for library preparation and pooling of libraries for sequencing on the NovaSeq 6000 sequencer. The workflow is optimized for minimal human touchpoints, with barcode verification and guided robotic handling to reduce error rates (Figure 7).

A key innovation in the IGA workflow is the optimisation of the library preparation protocol to remove the need for post-library QC prior to sequencing while maintaining a tight distribution of coverage (Figure 5 Box F). Libraries were pooled in batches of 24 per S4 flow cell, with correction factors applied to balance index representation. Clarity LIMS generated pooling spreadsheets that guide robotic liquid handling systems to prepare libraries at optimal concentrations. Based on initial sequencing of approximately 20,000 samples, pooling volumes were further refined to improve yield uniformity. Samples falling short of yield thresholds were either re-sequenced (>20Gb shortfall) or undergo top-up sequencing (1-20Gb shortfall), depending on the extent of the shortfall. Clarity LIMS also safeguards against index clashes during re-pooling by cross-referencing indexed libraries.

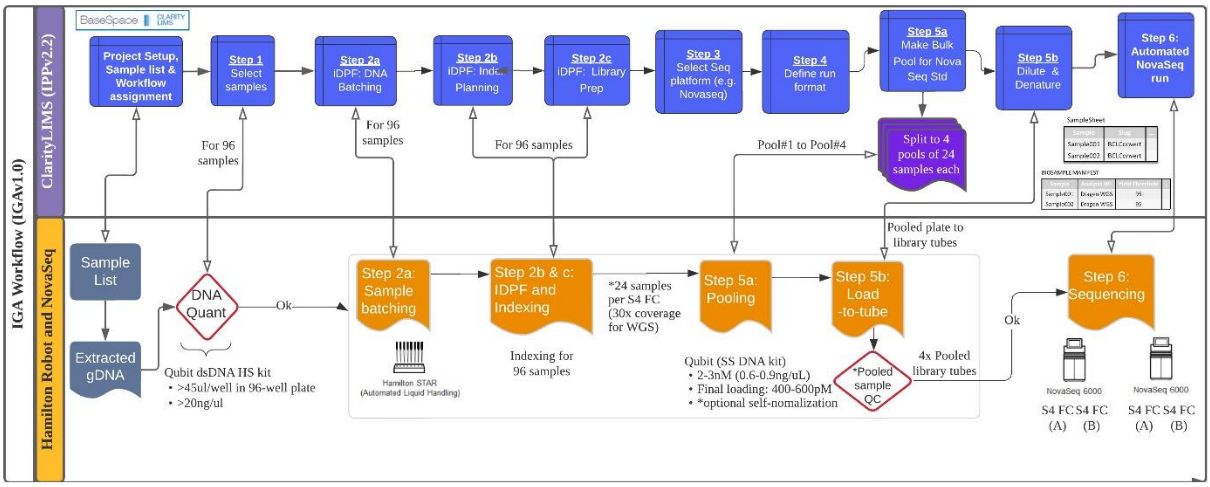

***Figure 6. Summary of the steps taken from library preparation to sequencing***

### ii) Illumina Data Analysis Module

The Data Analysis Module is a fully automated, cloud-based pipeline that processes raw sequencing data into high-quality genomic outputs delivered into a structured data repository in AWS (ap-se1) through a securely managed gateway (Figure 8). Data from the NovaSeq

6000 was streamed into Illumina Connected Analytics (ICA) via BaseSpace Sequence Hub. Primary analysis included demultiplexing and FASTQ conversion using BCL Convert (version 3.9.4), followed by sex prediction through alignment of the first 1 million reads of a sample FASTQ to the hg38 reference genome with BWA-MEM (version 0.7.17-r1188) and determining the ratio of reads mapping to chrX and chrY using a custom script. Secondary analysis was performed using DRAGEN (version 3.7.8), encompassing read alignment, variant calling, QC reporting, followed by variant annotation with Nirvana (version 3.9.1) and multiQC (version 1.9) report generation. Each analytical step was gated by predefined quality metrics (Table 3), and failures trigger automated alerts and manual review through the dedicated SRC (Supplementary Note 6: Operational Considerations in the PRECISE-SG100K Genomic Engine). In the case where samples need to be re library prepped and sequenced, only new data was retained and analysed with DRAGEN. In the case of top-up sequencing, FASTQ files from first pass and top up sequencing were aggregated prior to secondary analysis. Data delivery was triggered through an automated system when a sample met the agreed quality metrics after DRAGEN analysis or delivery was triggered manually following review and data acceptance.

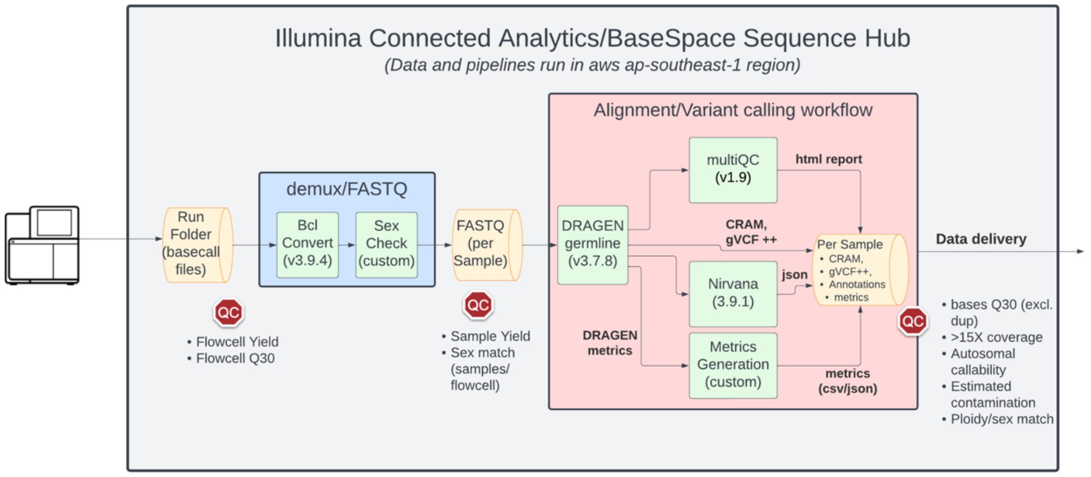

*__Figure 7. Data analysis and production workflow and pipelines.__ Data from the instrument was uploaded into the ICA cloud software via the BSSH software integration with NovaSeq 6000 instrument. Two analytics workflows were run on the data i) demultiplexing of raw data (bcl) into individual sample FASTQ files plus calculation of the predicted sex ii) processing of FASTQ files through a workflow that performed germline analysis (read alignment and variant calling) with DRAGEN 3.7.8, followed by generation of per-sample multiQC html report, annotation of variants with Nirvana and custom metrics aggregation. Each step was gated by specific QC parameters (defined in Table 3).*

#### iii) Illumina Sample and Data Management Dashboard

To ensure full traceability/auditability of the workflow, the project designed a simple but flexible data capture system in the Illumina Genomics Infrastructure. For every step of the process, we captured metadata related to input parameters, output metadata and within process updates (where applicable). All data was captured into an Amazon Web Services (AWS) DynamoDB table in a consistent structured format (Table 7) using AWS Simple Notification Service (SNS) messaging from Clarity LIMS and ICA (Figure 9). In total each sample had at least 96 process updates which captured name, process step, process stage, state (in progress, succeed, fail), timestamp and step metadata (structured in XML or json format) covering the entire lifecycle of the sample from submission through to delivery and data deletion. All operational management tools (see below) utilize data stored and captured in the main DynamoDB table.

***Table 7. Database structure used for tracking samples in the Illumina Data Generation Infrastructure.***

| **ID** | **Sample submission ID** |
|---|---|
| MajorStep | Tier 1 process step |
| StepName | Tier 2 subprocess step |
| Status | Started, succeeded, failed |
| Origin | Software of origin (Clarity, BSSH, ICA) |
| TimeStamp | Time when message was sent |
| AdditionalInfo | Json or xml structured additional data for the step (if applicable) |

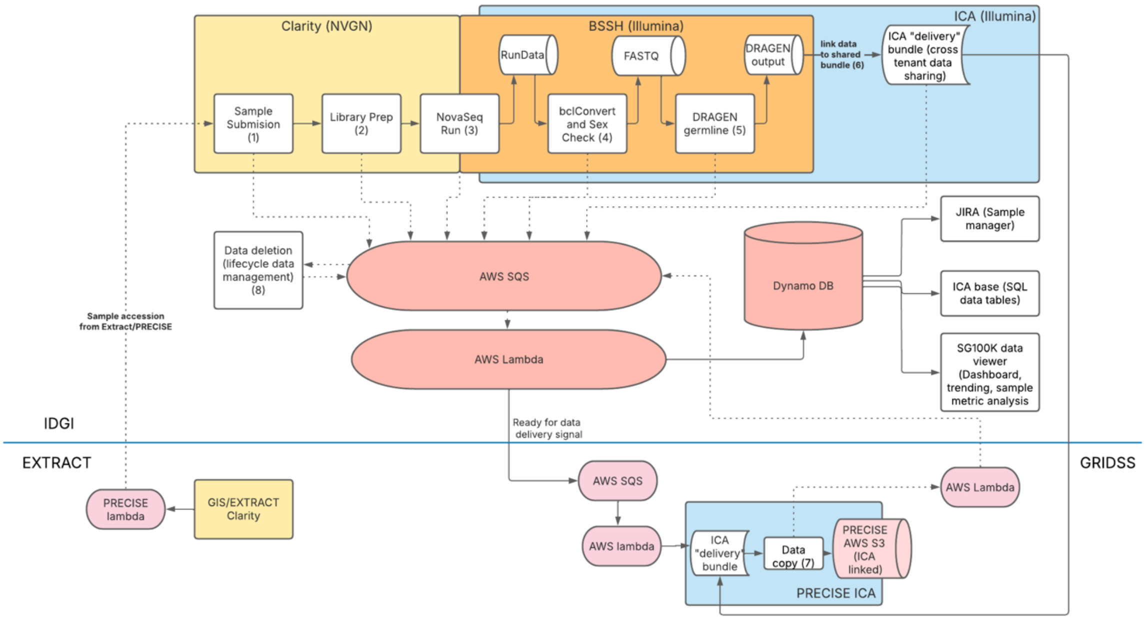

***Figure 8. Sample and process tracking for the Illumina Data Generation Infrastructure.*** *Each step in the process (numbered 1-8) from sample accession through to data delivery and data lifecycle management and deletion are captured in a central DynamoDB table. Wet lab processing steps (sample receipt to sequencing on NovaSeq 6000) are captured in Clarity LIMS (yellow), whereas data analysis and data delivery steps are managed with BSSH/ICA (Orange/Blue). Each step will send out a real-time SQS message (dotted line) which is captured by AWS SQS, and then an AWS Lambda will process the message and insert into the DynamoDB table in a consistent format. Once data generation has completed, an ICA bundle is used to facilitate data sharing, which involves a copy of the data into an ICA-linked and PRECISE managed AWS S3 bucket. Tools used for sample management (JIRA), data mining (SQL tables managed in the Base module of ICA) and sample monitoring, trending and plotting (SG100K viewer), ingest all data from the DynamoDB table.*

Operational monitoring is supported by a suite of tools that enable high-resolution oversight of sample progression. SQL tables within ICA's Base module allow complex querying of metadata, while a custom dashboard built with Python and Plotly provides daily operational insights and trend analysis. Additionally, Jira Cloud was used for sample-level tracking, with each sample assigned a ticket that is updated across 24 workflow stages. This system enabled rapid identification of QC failures and supported coordinated decision-making for reprocessing actions such as top-up sequencing or re-library preparation.

### Supplementary Note 5. Genome Research Informatics & Data Science - Informatics and data stewardship

#### Introduction

Informatics and data stewardship are managed by the Genome Research Informatics & Data Science (GRIDS) platform which deployed a secure environment for the processing, storage, and analysis of genomic data. GRIDS bridges the data production and downstream analysis.

Sample-level provenance, and data auditability were maintained through a Laboratory Information Management System, tailored to project-specific workflows, and complemented by purpose-built web-services for tracking of sample status from collection to analysis, linking to associated metadata, run performance metrics, and status logs.

#### PRECISE end-to-end Sample Tracking System

GRIDS built the PRECISE Sample Tracking System (STS) with the goal to record and monitor metadata, status and quality metrics associated with each sample flowing between the different modules of the PRECISE-SG100K Genomic Engine. The STS communicated with each of the parties and centralized relevant information.

#### Laboratory Information Management System (LIMS)

Samples' intake and DNA extraction module, termed EXTRACT, was supported by the Clarity Laboratory Information Management System (CLIMS) to handle SG100K sample metadata and laboratory sample tracking as well as data integration across all instruments and sample plates used in the DNA module. The system also communicates with the STS module to assign unique identifiers to each derived samples generated through the workflow.

Three workflows dedicated to EXTRACT activities were developed within Clarity LIMS v5.4: i) Sample Acceptance workflow, ii) Extraction workflow and iii) Re-submission workflow.

Each workflow and the overall LIMS system was built with the aim to reduce human touchpoints and record all actions applied to a sample. This enabled multiple benefits including the reduction of human error, an improved traceability of the processes, a better data organization and a more efficient data management.

**Sample Accessioning**

PRECISE STS ensured that each sample created through the workflow was assigned a unique identifier. Bio-samples received by EXTRACT were identified by a label provided by the sample provider. Through the automated DNA extraction and normalization workflow, two new samples were generated, 1) a stock DNA, raw result of the extraction and 2) a normalized DNA, aliquot to be sent to local sequencing service provider. During the relevant EXTRACT CLIMS step, messages were sent to a dedicated STS API endpoint that records the pre-barcoded 96 well plate ID and the well location of each sample in a dedicated database and assigned a unique identifier (NPMID). The NPMID was sent back to CLIMS, recorded, and attached to all downstream processes affecting the sample.

**DNA Extraction and Normalization Capture**

Participant metadata, DNA extraction and DNA normalization processes and quality metrics were initially recorded by EXTRACT into CLIMS. PRECISE STS systematically captured progress of each sample in the workflow by triggering customized scripts at the end of each CLIMS step using the built-in automation feature. A structured message was then sent to AWS SNS. The message was received and recorded into a comma separated file (CSV) saved on AWS S3. Using the Base module of ICA, CSV files stored in S3 were automatically loaded in a dedicated table.

**Genomic Data Delivery Capture**

Once high-quality genomic data was deemed ready to deliver to PRECISE, Illumina Data Generation Infrastructure (IDGI) sent a notification to the GRIDS Module through AWS SNS/SQS. The message was first recorded into AWS DynamoDB and then a workflow was triggered, running within the Flow module of ICA. This workflow copied the data from ILMN managed storage to PRECISE managed storage ensuring data handover and ownership. Once the transfer was completed, integrity of the data was confirmed by checking the list of files transferred against a manifest included with the data. A notification of transfer complete was then recorded into DynamoDB and sent to Illumina Genomics Engine (Figure 10). In case of transfer failure, GRIDS and ILMN teams investigated the root cause, and a new notification was sent manually to trigger the data transfer pipeline.

Processes related to DNA sequencing, mapping and variant calling were initially recorded by ILMN into the IDGI. The relevant tables generated into the Base module of ICA were then shared with PRECISE through a “Bundle” that allowed for secure and controlled accessed data sharing between multiple data owners within the platform (Figure 10).

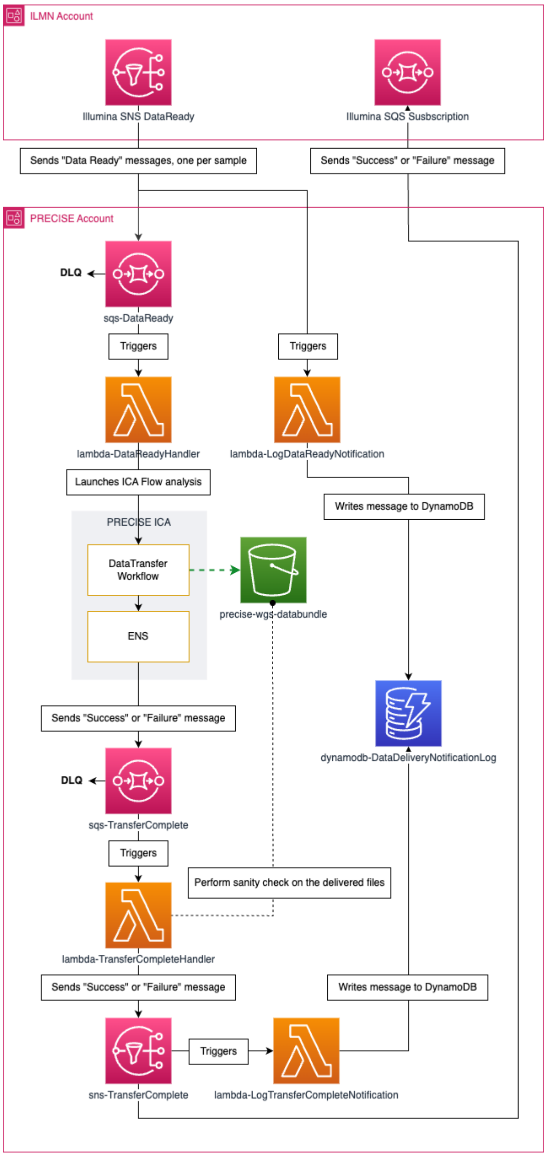

*Figure 9. Data bundle delivery workflow and automation*

**Sample Status Monitoring**

A monitoring application with controlled access limited to the NPM program stakeholders, was set up to allow the review of the sample status throughout the entirety of the PRECISE-SG100K Genomic Engine. In the backend, it queried the PRECISE STS database. The frontend used Plotly Dash framework. This application presented an overview of the current progress of the project, with count of samples at each stage of the workflow. It also allowed for the retrieval of detailed information of specific samples and all the associated events. Lastly, a dedicated section was focused on reviewing DNA & WGS QC metrics, trending specific metrics in real-time.

The development of this application, centralizing the information generated by the multiple structures and institutions involved in data generation, was instrumental for PRECISE management to review the progress of the project as a whole and critical for the operators to monitor the quality of production and gain insight of eventual issues.

### Supplementary Note 6: Operational Considerations in the PRECISE-SG100K Genomic Engine

#### Introduction

Beyond laboratory automation and informatics platforms, the success of the PRECISE-SG100K Genomic Engine depended on deliberate operational design. A national-scale program sequencing 100,000 whole genomes required structures for governance, communications, workforce planning, and continuous improvement that extended beyond technical workflows. Here we outline the operational frameworks that enabled consistent performance and on-time delivery.

#### Governance – Sample Review Clinic and Joint Steering Committee

Two tiers of governance underpinned day-to-day and strategic oversight.

- The Sample Review Clinic (SRC) convened every two weeks and served as a technical decision-making forum for samples that deviated from acceptance criteria. Using live GRIDS STS dashboards and Jira boards, stakeholders jointly determined corrective actions such as resequencing, top-up, or sample abandonment. This mechanism prevented backlogs and maintained uniform turnaround (Figure 1).
- The Joint Steering Committee (JSC) met every six months, comprising senior representatives from PRECISE, GIS, and ILMN. The JSC reviewed program milestones, resolved escalated issues, and monitored partner commitments. Its structured escalation pathway ensured that operational deviations could be rapidly surfaced and addressed at leadership level.

#### Communications and Workforce Management

The Engine's multi-party nature required structured communication channels. Microsoft Teams was adopted as the central platform, complemented by a standing meeting cadence across workgroups. Biweekly operational syncs were held for DNA extraction, sequencing, informatics, and data management teams. These meetings standardized agendas and ensured transparency across institutions. Workforce planning was equally deliberate: each module (EXTRACT, IDGI, GRIDS) was staffed with appropriately resourced teams, surge capacity, and redundancy to de-risk bottlenecks and ensure continuity.

**Ramp-up and Continuous Improvement**

Operations commenced with a six-month “Stress Test” of ~3,000 samples, intentionally run at reduced throughput to expose process vulnerabilities. Findings informed modifications to DNA quantity requirements, WGS acceptance thresholds, and queuing logic for resequencing and top-up, reducing number of samples entering SRC 3-fold. Periodic checkpoint reviews at 15,000 and 50,000 genomes quantified improvements in turnaround time and reduction of samples routed to SRC. These iterative refinements enabled the Engine to reach optimal efficiency of steady-state throughput within nine months, generating >3,000 genomes per month with <1% of samples requiring SRC adjudication.

**Operational Monitoring and Data Transparency**

Real-time monitoring was achieved through a combination of commercial and bespoke tools. The PRECISE STS, ICA, and AWS web services generated a live, queryable record of sample status. A PRECISE-SG100K dashboard enabled day-to-day tracking of throughput and key performance metrics, while Jira tickets provided sample-level audit trails. This infrastructure not only improved troubleshooting but also created institutional memory for continuous optimization.

**Cross-Partner Operations and Public-Private Collaboration**

A distinctive feature of the Genomic Engine was the deliberate co-development between public and private partners. Co-location of industry and research teams allowed real-time troubleshooting, while formal agreements on cost-sharing, intellectual property, and knowledge transfer aligned incentives across parties. Industry partners contributed not only technology platforms but also logistics expertise, training, and capacity planning, accelerating scale-up to national throughput.

**Key Takeaways / Considerations (summarized in Table 8)**

- Operational scaffolding was essential to transform technical workflows into a reproducible national-scale system.
- Two-tiered governance (SRC, JSC) provided complementary technical and strategic oversight.
- Structured communication channels and live dashboards minimized information asymmetry across partners.
- Phased ramp-up and checkpoint reviews enabled continuous improvement without compromising data quality.
- Co-development with industry partners reduced infrastructure duplication and accelerated troubleshooting.
- Workforce planning and surge capacity mitigated risks from staffing fluctuations and equipment downtime.

*Table 8. Summary of Operational Structures and Mechanisms Supporting the PRECISE-SG100K Genomic Engine*

| Operational Element | Function | Mechanism | Outcome |
|---|---|---|---|
| Sample Review Clinic (SRC) | Technical decision forum | Fortnightly cross-partner meeting using dashboards and Jira | Rapid resolution of non-conforming samples; <1% routed to SRC at steady state |
| Joint Steering Committee (JSC) | Strategic oversight | Six-monthly leadership meetings | Escalation of unresolved issues; alignment on milestones and change controls |
| Workforce communication | Daily/weekly coordination | Microsoft Teams channels; biweekly syncs per module | Transparency across partners; reduced information asymmetry |
| Ramp-up strategy | Safe transition to scale | Stress Test (~3,000 samples), 15K & 50K checkpoints | Iterative refinement of acceptance criteria; improved turnaround and yield |
| Operational monitoring | Real-time tracking | PRECISE Sample Tracking System; ICA; AWS logs; SG100K dashboard | Live visibility of throughput, QC metrics, and trend analysis |
| Workforce planning | Capacity and resilience | Right-sized staffing; surge capacity; redundancy | Stable throughput despite turnover or equipment downtime |
| Public-private collaboration | Alignment and knowledge transfer | Co-location, cost-sharing, IP frameworks | Accelerated implementation; shared responsibility for outcomes |

### Supplementary Note 7. Outcome

Delivered samples had a median coverage of 38X, with a minimum of 26X. Despite a small number of samples of less than 30X, the threshold used is likely to have a minimal impact of variant calling, with Single Nucleotide Variation / Indel having little change with higher coverage. However, we do see increase pass filter structural variant calling (both deletions and insertions) correlating with sample coverage.

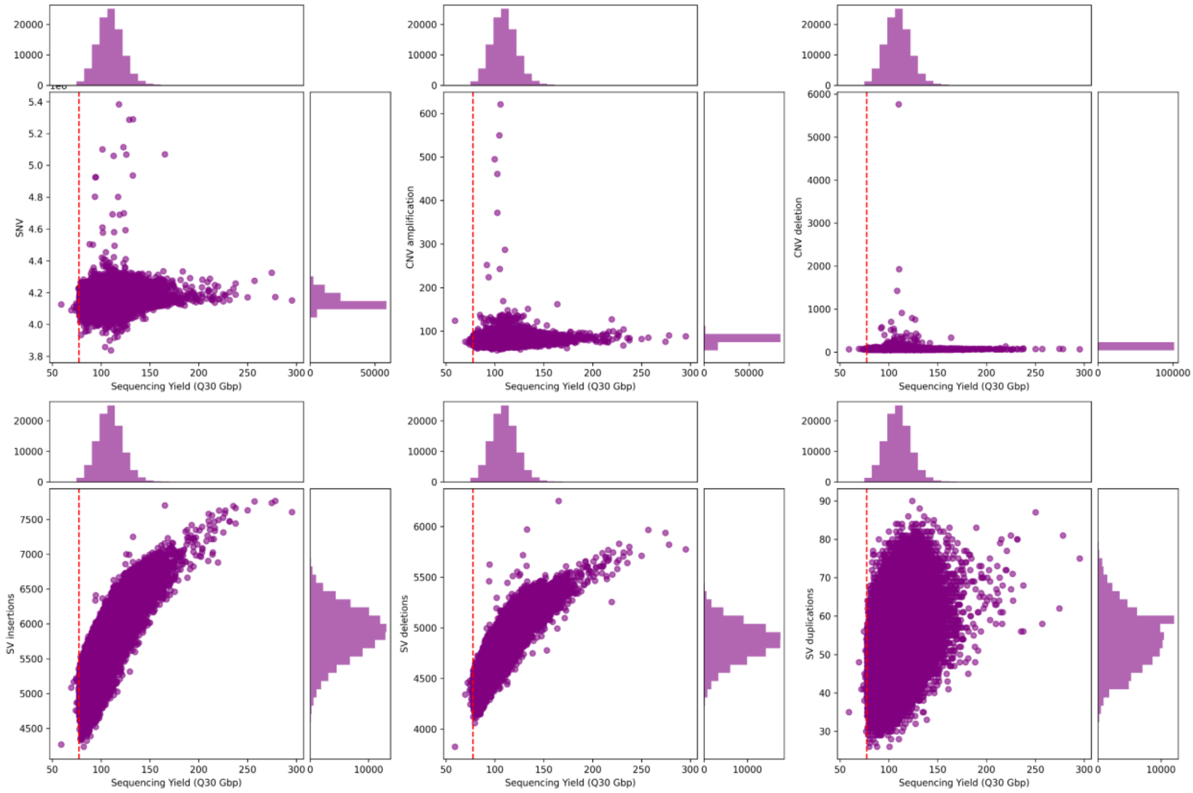

***Figure 10. SG100K-PRECISE Genomic Engine Sensitivity of Variant Calling against Yield for several classes of variant****. Sequencing yield measured as number of bases > Q30, removing duplicates and trimmed plotted against Single Nucleotide Variants; Copy Number Variant (CNV) amplification; CNV deletions; Structural Variant (SV) insertions; SV deletions and SV duplications*
